# Feeding and growth variations affect *δ*13C and *δ*15N budgets during ontogeny in a lepidopteran larva

**DOI:** 10.1101/2022.11.09.515573

**Authors:** Samuel Charberet, Annick Maria, David Siaussat, Isabelle Gounand, Jérôme Mathieu

## Abstract

Isotopes are widely used in ecology to study food webs and physiology. The fractionation observed between trophic levels in nitrogen and carbon isotopes, explained by isotopic biochemical selectivity, is subject to important within-trophic level variations, leading to imprecision in trophic level estimation. Understanding the drivers of these variations is thus important to improve the study of food webs. In this study, we characterized this variation by submitting *Spodoptera littoralis* larvae to a gradient of starvation levels, a factor that we hypothesized would change the trophic fractionation between individuals. The various growth rates that were induced from these starvation levels resulted in a ~ 1-1.5‰ within-trophic level variation of the trophic fractionation in both carbon and nitrogen, which is substantial compared to the 3-4‰classically associated with between-trophic levels variations. Hence starved animals sampled *in natura* may be ranked at a higher trophic level than they really are. We were able to gain an understanding of the effect of growth rate on isotopes fluxes between three easy-to-measure biological materials, food, organism and its wastes (frass), giving insight into physiological processes at play but also conveying helpful information to the sampling framework of field studies.

## Introduction

Stable isotopes are frequently used to understand fluxes of nutrients in ecosystems as well as trophic position and animal body condition (Post, 2002). The systematic differences in stable isotope levels between the resource and the tissue of a consumer - the trophic fractionation, here denoted Δ13C and Δ15N - are used to estimate the trophic level of consumers. It occurs because isotopes of different masses have slightly different kinetics during biochemical processes (i.e. respiration or absorption, see Fry, 2006). The 15N level of the consumer is usually increased by 3-4‰ relative to its resource because animals retain 15N preferentially over 14N (Martinez del Rio et al., 2009). Carbon fractionation, on the other hand, might vary in a population due to differences in the abundance of *de novo* synthesized lipids in the consumer’s body (Melzer and Schmidt, 1987). However, the within-trophic level variability of trophic fractionation sometimes impedes accurate trophic level estimation (Martinez del Rio et al., 2009). Understanding the drivers of these variations is crucial to improve our estimations.

Most of the proposed mechanisms to explain Δ15N variation involve diet protein quality and metabolism (Starck, Wang, et al., 2005). However, nutritional status, determined by the resource availability in the environment (Doi et al., 2017, Trochine et al., 2019), can influence trophic fractionation. Physiological responses to nutritional stress involve adjustments in digestion, reserve utilization and metabolic rate. As these processes change in rate (see fig.1.a. and c.), biochemical processes that determine absorption, respiration and excretion, also change, therefore impacting Δ15N and Δ13C. Total food restriction, which causes weight loss (corresponding to negative growth rates in fig.1.b. and d.), has the overall tendency to increase heavy isotopes content (15N and 13C), leading to an overestimation of the trophic level (see Adams and Sterner, 2000; Boag et al., 2006; Gorokhova and Hansson, 1999; Haubert et al., 2005; McCue, 2008; Oelbermann and Scheu, 2002; Olive et al., 2003; O. Schmidt et al., 1999). But more rarely have the effect of various feeding levels been considered, with no convincing conclusion to this day (Hertz et al., 2015). To improve the estimation of trophic levels by including these mechanisms, we need a detailed understanding of the relationship between variation in nutritional status and trophic fractionation.

**Figure 1:**
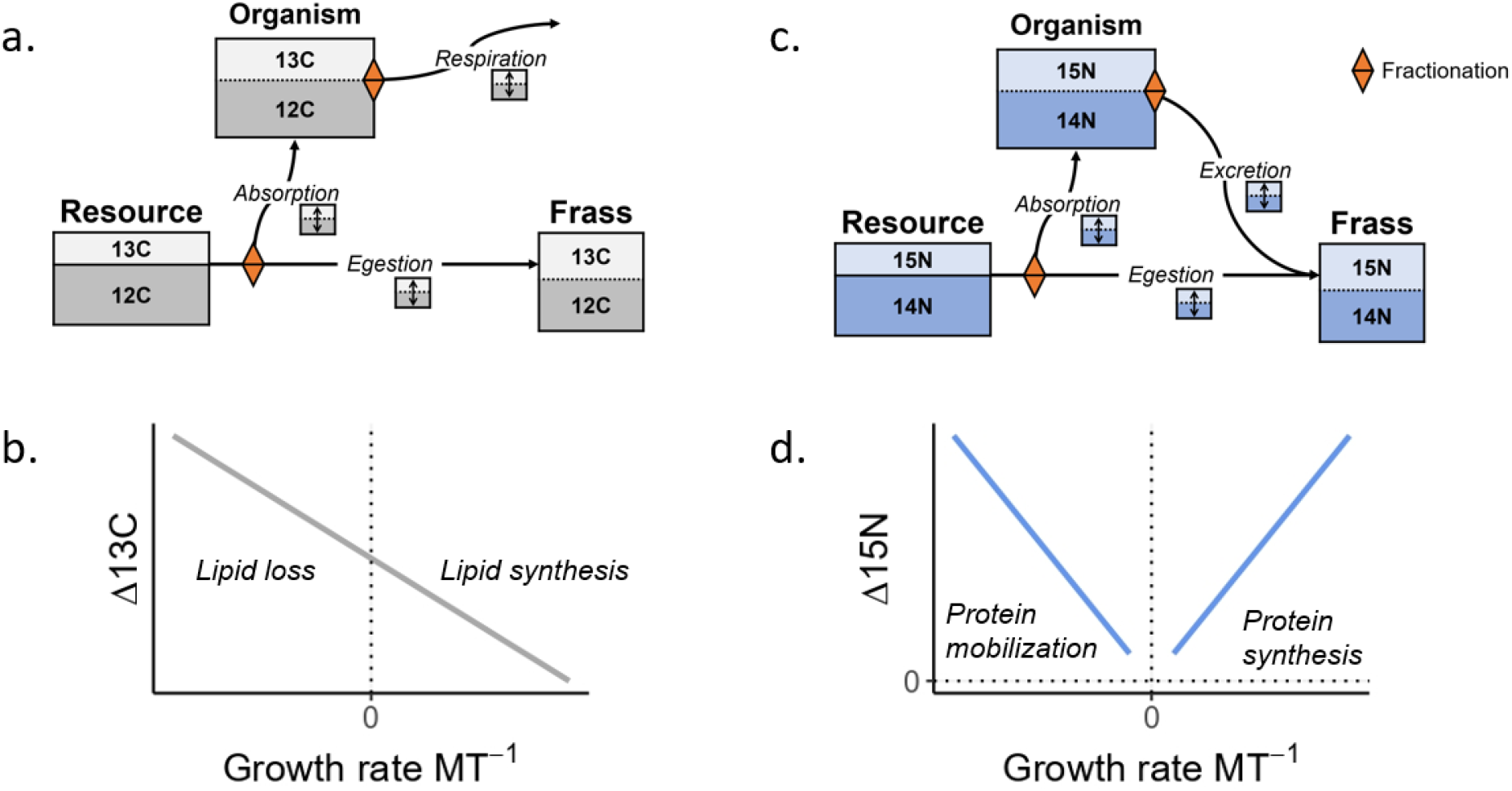
Isotopes routing and main hypotheses. a. & c. The three analyzed matrices, their relative (not-to-scale) content of isotopes, fluxes between them, as well as nodes where fractionation can occur (diamonds). The exact proportion of isotopes is not intended to represent reality faithfully but rather to illustrate the dynamical aspect of trophic fractionation. b. & d. Hypothesized relationship between trophic fractionation and growth rate for nitrogen and carbon. a. Most carbon is lost through either respiration or egestion and marginally through excretion. c. On the contrary, nitrogen is solely dropped through either egestion or excretion, with the impossibility of distinguishing their contribution only based on frass analysis. b. The hypothesized relationship between Δ13C and growth rate, measured as mass gained per unit of time *MT*^-1^. We expect a negative relationship because of the increasing proportion of 13C-poor *de novo* synthesized lipids, thus modulating the respiration fractionation. d. Hypothesized relationship between Δ15N and growth rate. High growth rates should increase protein synthesis and breakdown rates, which retain preferentially 15N, and very low intake rates (weight loss) should increase protein catabolism, also increasing Δ15N, both playing on the excretion fractionation.

Across this gradient in nutritional status, an important threshold is the maintenance feeding level (zero growth rate in fig.1.b. and d.). Below the feeding level required for maintenance, body mass decreases, and adaptations in lipids and proteins metabolism are triggered. Lipids typically contain proportionally less 13C than proteins and carbohydrates (DeNiro and Epstein, 1977; McConnaughey and McRoy, 1979). A shrinkage of body lipid content should thus result in an increase in Δ13C compared to high feeding levels (Gaye-Siessegger et al., 2004), where the organism might be able to accumulate 13C-poor lipid reserves, therefore decreasing Δ13C (fig.1.a.).

Regarding nitrogen, low feeding levels are classically associated with an increase in Δ15N due to asymmetrical isotopic routing during protein mobilization for energetic catabolism (Hatch, 2012, see fig.1.c.). But high feeding levels, which are often accompanied by high growth rates, can also be accompanied by an increase in Δ15N (Sick et al., 1997, Focken, 2001). Indeed, due to an increase in protein synthesis and breakdown rates when the animal is growing fast, removal of 14N is intensified, thus enriching the consumer in 15N and increasing Δ15N (fig.1.c.). As a result, both very low and very high intake rates might increase Δ15N, but due to different processes, protein mobilization at low intake rates in a weight loss context and protein synthesis and breakdown rates at high intake rates in a growth context.

Moreover, as the gut filling level decreases with underfeeding, the food passage time increases and the biochemical conditions in the gut change. This change in temporal and chemical conditions might alter the isotopic fractionation right from the absorption stage (H.-L. Schmidt et al., 2015). The relative decrease in the concentration of food in the near-empty gut might increase the enzymes’ accessibility and, in turn, the absorption of heavy isotopes. As a whole, trophic fractionation should depend on both nutritional status and body mass dynamics (Sears et al., 2009, Williams et al., 2007; see also Hatch, 2012 for a review), but these effects remain poorly investigated, especially in varying feeding levels (see Gaye-Siessegger et al., 2007).

Elucidating how the nutritional status modifies isotopic fractionation in a growing organism could shed light on the within-trophic level variability of the estimated trophic level and should be of interest for field studies as well. We conducted a feeding level experiment during the larval development of the cotton leaf worm *Spodoptera littoralis,* including severe food restriction, three intermediate restriction levels, and an *ad libitum* level, which corresponded to a range of growth rates. We assessed 15N and 13C budgets, measuring isotopic fractionations between food, body and frass (excretion + egestion).

More specifically, we wanted to test the hypotheses that:

1. Δ15N should increase at negative growth rates due to protein catabolism during weight loss and increase at positive growth rates due to faster protein synthesis and oxidation. Around maintenance level, as these two processes slow, Δ15N should decrease. Overall we should thus obtain a V-shaped relationship between Δ15N and growth rate (fig.1.d.).
2. Δ13C should decrease with growth rate because of the accumulation of 13C-poor lipid stores (fig.1.c.).
3. The relative absorption of 13C might increase at low feeding levels as both gut passage time and digestion efficiency increase.

## Material and methods

### Study system

*S. littoralis* larvae from a laboratory strain were reared on a semi-artificial diet for the total duration of the experiment. We provide the detailed food composition in Appendix 1, table 1. The climate chamber was set at 23 °C, 60–70% relative humidity, and a 16:8 light/dark cycle (Hinks and Byers, 1976). In these rearing conditions and with continuous access to food, the larvae go through 7 instars before entering metamorphosis (chrysalid stage). To enable proper mass balance calculation and prevent cannibalism, we isolated the 400 larvae intended for the experiments at the 6th instar in individual 30 mL circular polypropylene boxes. We provided them *ad libitum* food until 6th moult completion (start of the 7th instar).

**Table 1:** Summary of linear models describing the influence of growth rate and mass-specific ingestion rate (MSIR) on the trophic fractionation (Δ), and carbon isotope absorption efficiencies ratio (C IAER), respectively.

| Equation | n | $R^2$ | F | p-value |
| --- | --- | --- | --- | --- |
| $\Delta^{13}\text{C} = -0.0032 \times GR + -1.7$ | 92 | 0.35 | 48 | $p < 0.01$ |
| $\Delta^{15}\text{N} = 0.0054 \times GR + 0.32$ | 92 | 0.53 | 100 | $p < 0.01$ |
| $\text{C IAER} = -0.65 \times \text{MSIR} - 0.97$ | 100 | 0.28 | 38 | $p < 0.01$ |

### Experimental design

We randomly assigned each of the 400 7th instar larvae to one of five food provision levels for the duration of the experiment. Food was kept the same as before the start of the feeding level experiment. The food intake level was fixed to either 120, 240, 360, 480 or 900 mg of food per day (fw), depending on the larva. We had beforehand estimated the average maximal individual intake rate for 7th instar larvae and obtained 595 ± 43 mg/day. There were 80 individuals for each tested food intake level. We conducted this study over 10 weeks (10 temporal blocks), performing the experiment with 40 individuals each week, 8 for each food intake level. Individual measurements and sample collections took place over two or three days depending on whether the larva pre-pupation occurred on the third day of the 7th instar (in which case measures were taken during 2 days) or later (in which case measures were taken during 3 days).

### Experimental workflow

During the experiment, each larva was given the defined amount of freshly prepared food and weighed every day. Food subsamples were taken at every food preparation for subsequent chemical analysis. We collected and weighed daily food leftovers and frass produced by each larva to assess the actual intake and egestion rates. Food leftovers and frass were quickly stored at −20 °C, and later dried for 72 hours at 60 °C in an oven, to measure their dry mass. On the third day, half the larvae were quickly stored at −20 °C, dried for 72 hours at 60 °C in an oven, and their dry mass was measured. The other half of the individuals was left in the rearing chambers to later investigate the effect of food restriction on mortality, emergence success and body mass (not analyzed here).

### Chemical analyses

Chemical analyses required that we pooled samples to obtain enough analyzable material. Hence groups of 4 caterpillars reared over the same week and on the same feeding level were composed, 2 that were pooled together for chemical analysis, and 2 that were left alive until emergence. The analyzed frass was a pooled sample of all 4 individuals.

All samples - food, larvae, and frass - were ground to a fine powder using a mill. Total carbon, total nitrogen, as well as *δ*13C and *δ*15N were measured using an elemental analyser coupled to a mass-spectrometer (Flash HT - Delta V Advantage, ThermoFisher). We checked for measurement errors using aromatic polyimide (EMA-P2) as standard.

### Starvation proxy and isotopic data

Intake rate alone does not accurately represent the nutritional status. Rather, it depends on the balance between intake and requirements, the latter largely depending on body mass. We, therefore, used mass-specific ingestion rate (MSIR) as an indicator of nutritional status. Low values of mass-specific ingestion rate define intense starvation, whereas high values of mass-specific ingestion rate represent sufficient intake.

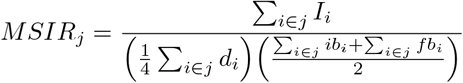

with *I_i_* the total fresh mass of food ingested by the individual *i* of the group *j* over the course of the 7th instar, *d_i_* the number of days spent in 7th instar by individual *i*, *ib_i_* the initial body mass of individual *i*, and *fb_i_* the final body mass of individual *i*.

Pee Dee Belemnite (PDB) and atmospheric nitrogen were used as standards for *δ*13C and *δ*15N, respectively. Isotopic data for sample *s* are reported using delta notation:

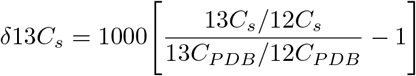

and

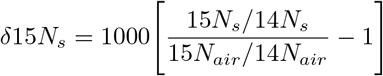

The trophic fractionation, *i.e.* the difference in *δ*13C or *δ*15N between larvae and food, was computed as follows:

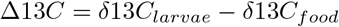

and

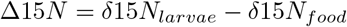

We computed the ratio of absorption efficiencies between the two carbon isotopes (thereafter C IAER) to characterize how isotopes are differentially absorbed. This metric characterizes the absorption process, which is one of the two fluxes, along with respiration, determining carbon trophic fractionation (fig.1.a.). We did not compute this metric for nitrogen because, unlike carbon, nitrogen excretion products also end up in insect frass, and it is, therefore, impossible to disentangle absorption from excretion effects using this metric (fig.1.c.). Moreover, as samples are heated during drying, some ammonium might volatilize, biasing the mass balance (Harrison, 1995).

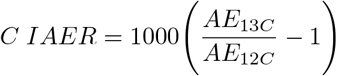

with *AE_i_* the proportion of ingested isotope which is assimilated, and not egested/excreted, over the 7th instar, given in % dry weight:

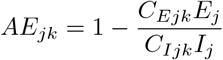

with *C_Ejk_* the proportion of isotope *k* in the frass of the group *j*, *E_j_* the summed mass of frass produced by the four larvae of the group *j* and with *C_Ijk_* the proportion of isotope *k* in the food of the group *j*, *I_j_* the summed mass of food ingested by the four larvae of the group *j*. Please refer to Appendix 2 for a detailed calculation of *C_Ejk_* and *C_Ijk_*.

### Statistics

To test the effect of starvation and subsequent variation in growth rate (GR) on the trophic fractionation and relative carbon isotope absorptions, we used linear regressions. We chose to test the effect of growth rates on Δ15N and Δ13C, and the effect of mass-specific intake rate on C IAER. Details on modelling choices are provided in Appendix 3.

## Results

Despite strong starvation conditions, we were not able to force negative growth rate (fig.2.a.). We were therefore unable to test the relationships between trophic fractionation - Δ13C and Δ15N - and growth rates for negative growth rates. Here, we describe these relationships for positive growth rates only.

**Figure 2:**
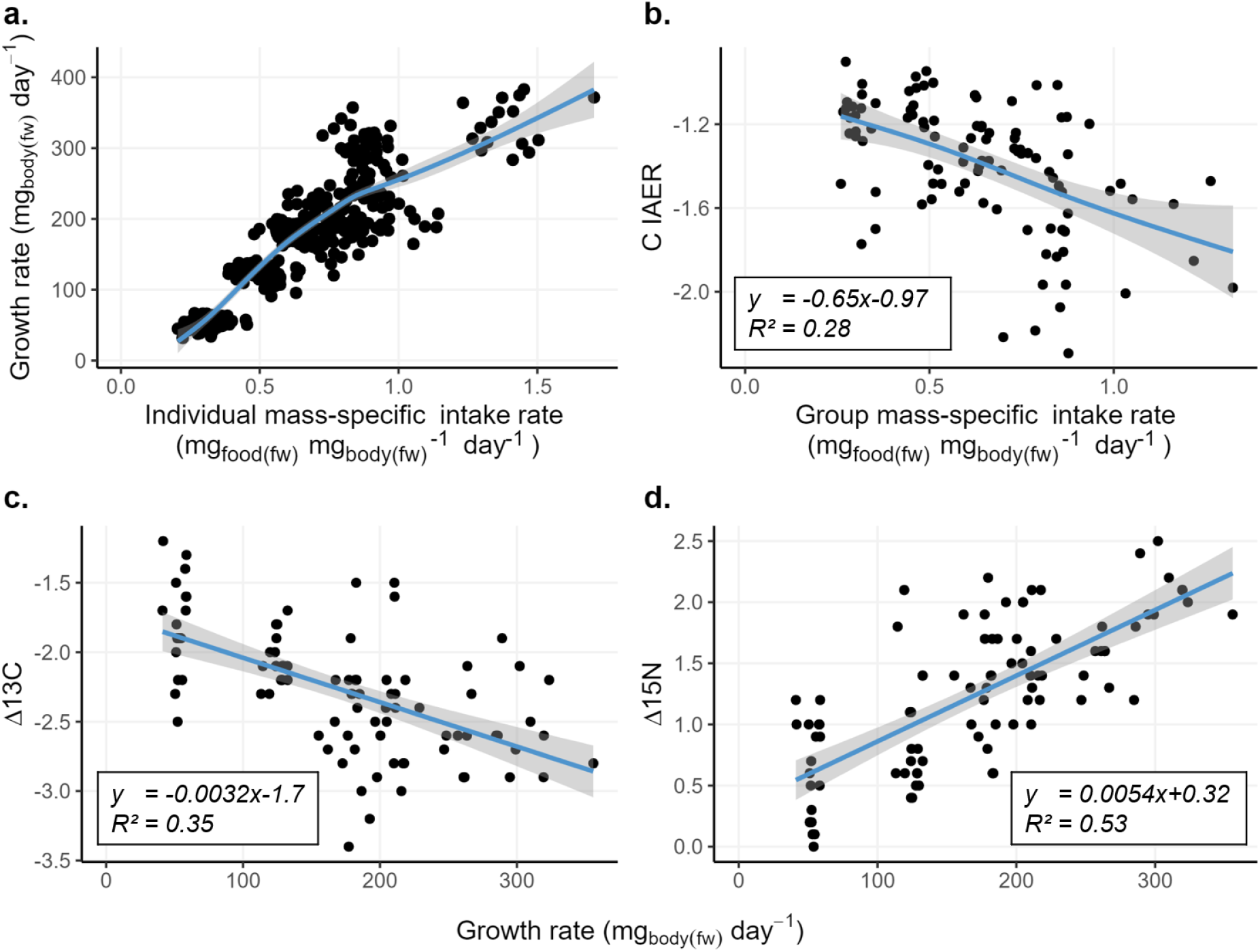
Growth and isotopic analyses. a. Individual growth rate as a function of mass-specific intake rate. The fitted curve is a generalized additive model. b. Carbon isotope absorption efficiencies ratio (IAER) as a function of mass-specific intake rate measured at the level of a group of 4 caterpillars, hence the term “group”. c. Carbon trophic (Δ13C) fractionation as a function of growth rate. d. Nitrogen trophic fractionation (Δ15N) as a function of growth rate.

### Trophic fractionation

As expected, larvae were always richer in 15N than the food they ate (Δ15N>0 for all larvae, see fig.2.d.). There was a clear positive correlation between Δ15N and (positive) growth rate (*F* = 100, *p* < 0.01, *R*^2^ = 0.53 ; table 1.) in accordance with our hypothesis (right side of the graph in fig.1.d). As for carbon, larvae were always poorer in 13C than their food (Δ13*C* < 0 for all larvae, fig.2.c.), and this difference was exacerbated by a growth rate increase (*F* = 48, *p* < 0.01, *R*^2^ = 0.35 ; table 1.), also in accordance with our hypothesis (fig.1.c.). In both cases, the Δ13C and Δ15N spanned over a range of ~ 2.5 ‰, of which 1 ‰ in the case of carbon, and 1.5 ‰ can be fully attributed to growth rate variation. These variations are substantial vis-à-vis the one classically attributed to a one trophic level shift (3-4 ‰),

### Isotope absorption efficiencies ratio (IAER)

The relative absorption of 12C and 13C depended on the mass-specific intake rate. 12C was systematically better absorbed than 13C, and this effect increased with feeding level (*R*^2^ = 0.28, *F* = 38, *p* < 0.01, fig.2.b.).

## Discussion

In agreement with our prediction, Δ15N increases with growth rate, up to 1.5 ‰, which is substantial compared to differences typically associated with trophic fractionation (3-4 ‰). Our results agree with previous work showing that Δ15N is sensitive to growth, at least in some tissues, as highlighted by Sick et al., 1997. We show that food limitation does not always increase Δ15N but rather depends on the underfeeding intensity and whether underfeeding is concurrent with growth. This contrasts with the classic view that Δ15N should increase in starved individuals owing to protein depletion for energetic requirements. At least two studies suggested that this increase in Δ15N at high growth rates could be due to higher rates of deamination and protein synthesis at higher intake rates (Sick et al., 1997, Focken, 2001). Combining both predictions leads to a more comprehensive view of the effect of feeding level on nitrogen trophic fractionation. Despite very low intake rates, down to 10% of *ad libitum* levels, no weight loss was observed in our experiment, leaving the complete shape of the relationship between Δ15N and growth rate only speculative, although the fact that most studies show an increase of Δ15N with starvation intensity or fasting duration in a negative growth context, whereas we find the contrary for positive growth rates, suggest such a relationship (Del Rio and Wolf, 2005). But whether a V-shaped relation can arise or not requires further investigation.

On the other hand, Δ13C decreased with growth rate and intake level, which is consistent with previous findings (Doi et al., 2017). This is likely due to the possibility of constituting 13C-poor lipid reserves at high growth rates (DeNiro and Epstein, 1977, McConnaughey and McRoy, 1979). To conclude, both Δ13C and Δ15N were affected by feeding level and growth rate. This shows that when assessing trophic levels using isotopic data, the nutritional status of the individual can bias the estimate. Despite being hard to estimate without destructive measurements, at least severe starvation and underfeeding might be detectable through environmental conditions. Physiologically, the nutritional state at which an individual grows can be assessed through age-size comparison, sclerochronology if applicable (Castanet, 1994), or biochemical indicators (e.g. ketone bodies, Chowdhury et al., 2014; Shah and Bailey, 1976).

Diet indicators are also prone to estimate error owing to variable nutritional status. The carbon isotopic signature of herbivores is sometimes used to estimate if their diet is composed primarily of C4 plants, rich in 13C (12 to20 ‰), or of the 13C-poorer C3 plants (25 to32 ‰), O’Leary, 1981). Elevated *δ*13C values in the consumer can hence indicate a predominance of C4 plants in the diet. The proportion of C3 in the diet of insects has sometimes been inferred through this tool. It is not clear whether the intensity of isotopic fractionation due to starvation could change as a result of the difference between C4 or C3-based diets, the present case being an example of an artificial diet containing both. Still, starvation is likely to lead to overestimates of the C4 fraction, although not by much (around 10% based on fig. 3 in Fry et al., 1978).

The mass budget of heavy and light isotopes revealed that 12C was more easily absorbed than 13C, which is consistent with the observation of a negative Δ13C. But as the intake rate decreases, 13C is better absorbed compared to well-fed animals. This indicates that the biochemical environment of the gut varies with intake level, with effects on the processes of digestion and absorption. Moreover, we can also conclude that the respiration fractionation either is negligible compared to the one associated with absorption or that it further decreases the amount of 13C in the organism. But the biochemical origin of this modulation of 13C absorption is unclear. It could be due to longer gut passage time, or to increased food enzymatic availability at low gut filling levels. Our results reveal that the within-trophic level differences in trophic fractionation imputable to nutritional status (1-1.5 ‰) are substantial compared to differences typically associated with trophic level changes (3-4 ‰). Hence assessing trophic levels *in natura* using isotopic analysis requires caution, especially if the community is perturbed and might be subject to nutritional stress. With the changes in frequency and intensity of drought episodes, one should be cautious to these potential biases in isotopic trophic ecology.

## Contribution

S.C., A.M., J.M., D.S. and I.G. designed the study. S.C. ran the experiment, did the analyses, and wrote the first draft of that manuscript. All authors contributed to the final version of the manuscript.

## Acknowledgements

The authors wish to thank Anabelle Fuentes and Philippe Couzi for their contribution to rearing and Magloire Mandeng-Yogo for his support to chemical analysis.

## Data, script and code availability

Data and code are available at DOI 10.5281/zenodo.7327343.

## Appendix

### 1 Food ingredients

**Table 1:** Composition of the feed distributed to larvae, expressed as % mass/mass.

| Ingredient | % m/m |
| --- | --- |
| Deionized water | 76.7 |
| Soja meal | 6.79 |
| Corn flour | 6.79 |
| Germalyne | 3.40 |
| Yeast | 2.55 |
| Agar | 1.20 |
| Casein | 7.19E-01 |
| D-Glucose | 6.01E-01 |
| Ascorbic acid | 5.10E-01 |
| Benzoic acid | 2.69E-01 |
| Linseed oil | 1.92E-01 |
| Nipagin | 1.16E-01 |
| Choline chloride | 5.41E-02 |
| Formaldehyde | 3.60E-02 |
| Alpha-Tocopheryl acetate | 1.59E-02 |
| Actitetra (Oxytetracycline 50%) | 9.59E-03 |
| Ampicillin sodium salt | 7.19E-03 |
| Myo-inositol | 3.61E-03 |
| Nicotinic acid | 3.21E-03 |
| Menadione | 1.62E-03 |
| Retinyl acetate | 1.30E-03 |
| Riboflavin | 7.21E-04 |
| Pyridoxine | 7.21E-04 |
| Thiamine hydrochloride | 7.21E-04 |
| Ergocalciferol | 9.02E-05 |
| Folic acid | 6.49E-05 |
| Biotin | 1.44E-05 |
| Cobalamin | 9.74E-07 |

### 2 Isotope absorption efficiencies ratio (IAER) and *C_Ejk_* / *C_Ijk_* calculation

Mass spectrometer usually directly gives isotopes ratio rather than isotopic content because usually, one of the isotopes has a low concentration.

Still, it is possible to compute the isotope content of egestion *C_Ejk_* and intake *C_Ijk_*.

For carbon, ignoring the very low concentration in unstable isotopes, we have that the total carbon content is equal to the sum of the content of each stable isotope. So that, for sample *s*:

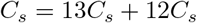

On the other hand the isotopic data are usually given in delta notation:

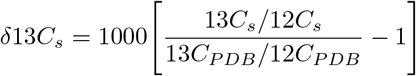

We have thus two unknowns, 13*C_s_* and 12*C_s_*, as well as two equations, enabling us to solve for the two isotopes content:

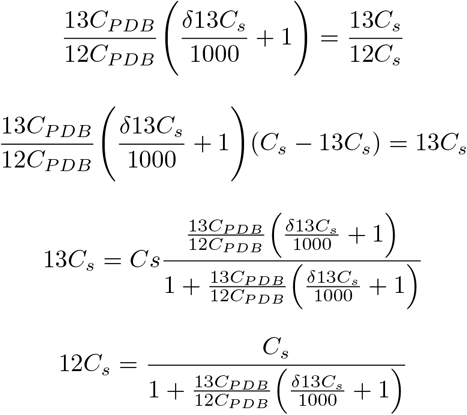

We have that 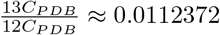, so, finally:

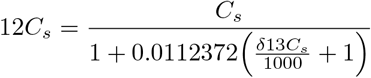

Using the isotopic content, we can compute the absorption efficiency of each isotope.

### 3 Justification of the choice of linear models

We predicted that growth experienced during a given period at a certain rate would affect the isotopic content of the organism, that is, taking the example of carbon, which also holds for nitrogen:

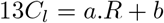

with 13*C_l_* the 13C content in the larva, *R* the growth rate, *a* and *b* some constant, we should have *δ*13*C* expressed as a function of growth rate *R* as follows:

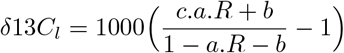

which is a hyperbolic function of *R* (*c* here is the standard isotopes ratio constant). We should thus expect non-linearity. However, as 13*C_l_* is very low, and making the approximation that for 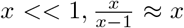 we can model this relation using a linear approach. We nevertheless tested for non-linearity by performing generalized additive models and examining the effective degree of freedom (edf).

**Table 2:** Generalized additive models results, with the sample size n, the effective degree of freedom (edf), p-value of the smooth term and the *R*^2^.

| GAM formula | n | EDF | p-value | $R^2$ |
| --- | --- | --- | --- | --- |
| $\Delta^{13}\text{C} \sim \text{s}(\text{GR})$ | 92 | 2.03 | <2e-16 | 0.368 |
| $\Delta^{15}\text{N} \sim \text{s}(\text{GR})$ | 92 | 1 | <2e-16 | 0.524 |
| $\text{C IAER} \sim \text{s}(\text{MSIR})$ | 100 | 1 | <2e-16 | 0.27 |

For the two trophic fractionations and C IAER, the EDF indicate a linear dependence, with EDF roughly between 1 and 2 (2). We therefore chose to use linear models.

